# Host age shapes virome abundance and diversity in birds

**DOI:** 10.1101/2020.08.23.263103

**Authors:** Michelle Wille, Mang Shi, Aeron C. Hurt, Marcel Klaassen, Edward C. Holmes

## Abstract

Host age influences the ecology of many microorganisms. This is evident in one-host – one virus systems, such as influenza A virus in Mallards, but also in community studies of parasites and microbiomes. We used a meta-transcriptomic approach to assess whether host age is associated with differences in the abundance and diversity of avian viromes. We used samples from cohabiting Ruddy Turnstones (*Arenaria interpres*) across three age groups, collected at two contrasting points in their annual migratory cycle. Overall, we identified 14 viruses that likely infect birds, 11 of which were novel, including members of the *Reoviridae*, *Astroviridae*, *Picornaviridae*, and *Phenuiviridae*. Strikingly, 12 of the viruses identified were from juvenile birds sampled in the first year of their life, compared to only two viruses in adult birds. Similarly, both viral abundance and alpha diversity were higher in juvenile than adult birds. Notably, time of sampling had no association with virome structure such that the migratory period may not play a major role in structuring avian viromes. As well as informing studies of virus ecology, that host age impacts viral assemblages is a critical consideration for the future surveillance of novel and emerging viruses.

## Introduction

Vertebrate animals of different ages also differ in their exposure and susceptibility to microorganisms [1]. The importance of host age for the epidemiology and ecology of microorganisms has been demonstrated by differences in the diversity and prevalence of parasites [2], bacteria [3, 4], and viruses [5–8] among hosts of different ages. In the case of wild birds, host age is associated with a higher prevalence of viruses in juvenile compared to adult birds. These include beak-and-feather disease virus in parrots [9], avian pox virus in albatross and tits [10, 11], avian avulavirus-1 in cormorants and waterfowl [12, 13], infectious bronchitis virus in poultry [14] and influenza A virus in wild waterfowl [8].

Despite a similar potential for exposure, viral susceptibility varies among adult and juvenile birds. This is believed to be due to an increase in the immune repertoire with age, with the age-related accumulation of immunity to influenza A virus in Mute swans (*Cygnus olor*) an informative case in point [15]. This increase of immunity in adults, coupled with juvenile immunologically naïve birds entering populations, may explain the significant increase in the prevalence of influenza A virus in juveniles compared to adults [7, 8], regardless of the time point of sampling [16–21]. This is evident when repeatedly sampling sentinel ducks, in which individuals have multiple influenza A virus infections in the first 6 months of age, compared to no or only short infections in the subsequent year of life, despite similar viral exposure [22]. In addition to varying susceptibility, exposure to viruses may similarly vary in wild birds. For example, in migratory birds such as shorebirds, adults and juveniles undertake southward migrations at different times, and in some species use markedly different routes, resulting in different virus exposure potential [23, 24].

As host age impacts the presence and prevalence of a range of specific viruses, we ask here whether host age is also associated with modulating communities of viruses carried by birds? To this end, we used the virome-scale data obtained from metagenomic next-generation sequencing as means to go beyond the “one-host, one virus” model and provide key data on the abundance and diversity of *all* the viruses present in a particular species. We used Ruddy Turnstones (*Arenaria interpres*) as a model species as they are known to be important reservoir hosts for viruses such as influenza A virus as well as for an abundance of other viruses [25, 26]. In the East Asian-Australasian Flyway, this species breeds in Arctic Siberia and following migration, spends its non-breeding period along the shorelines of Australia. Accordingly, we sampled individuals across three age groups at two different points in their annual cycle, just after migration, upon arrival in Australia and again shortly before departure to their Arctic breeding grounds. From these data we demonstrate that juvenile birds have significantly higher viral abundance and diversity compared to adult birds and that different time points in their migratory cycle had no apparent effect on virome strucutre.

## Methods

### Ethics statement

This research was conducted under approval of the Deakin University Animal Ethics Committee (permit numbers A113-2010 and B37-2013). Banding was performed under Australian Bird Banding Scheme permit (banding authority number 8001). Research permits were approved by the Department of Primary Industries, Parks, Water and Environment, Tasmania (permit number FA 13032).

### Sample collection

All birds were captured and sampled as part of a long-term study program on King Island, Tasmania, Australia (39°55’52”S, 143°51’02”E). Birds were captured by cannon netting at two time points: November 2014 following arrival from migration, and March 2015 prior to departure for migration to Siberia. Birds were aged using plumage characteristics and divided into three age categories: (i) 1o - juvenile birds hatched the previous breeding season, (ii) 2o – adolescent birds in their second year of life that overwintered on King Island (these birds could only be identified in November) and (iii) 3+ - adult birds, classified as those that are 3 years and older (Fig 1). A combination of oropharyngeal and cloacal samples were collected using a sterile tipped applicator and placed in virus transport media (VTM; Brain-heart infusion broth containing 2 × 106 U/L penicillin, 0.2 mg/ml streptomycin, 0.5 mg/ml gentamicin, 500 U/ml amphotericin B, Sigma).

**Figure 1.**
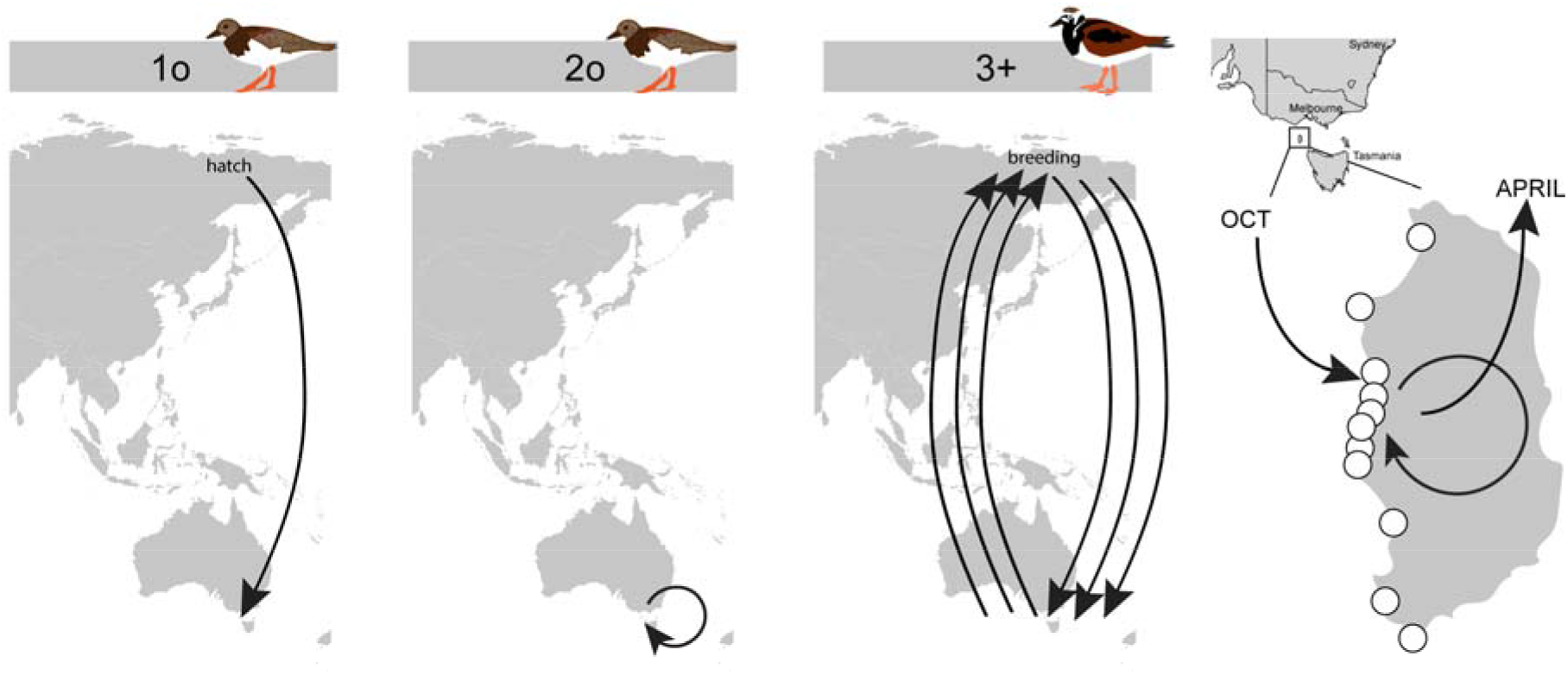
Sampling scheme. Birds occupied three age categories: juvenile (1o) birds were individuals hatched in Siberia that had performed a single migration leg to King Island; adolescent (2o) (that could only be identified based on their plumage in November) were individuals who had overwintered in Australia; and adult (3+) birds that perform an annual migration to Siberia and back to breed each year. Ruddy Turnstones arrive on King Island in October and can be found on rocky shores of the western coastline of the island throughout the Australian summer period. Small groups of ~50 birds with high site fidelity occupy beaches, with surveyed beaches denoted in small white circles. In autumn, following a period of intense weight gain, adult birds depart on migration to their breeding areas in March/April. Samples used in this study were collected in November 2014 and March 2015.

### RNA library construction and sequencing

RNA library construction, sequencing and RNA virus discovery was carried out as described previously [25, 27]. Briefly, RNA was extracted from swabs using the MagMax *mir*Vana™ Total RNA isolation Kit (ThermoFisher Scientific) on the KingFisher™ Flex Purification System (ThermoFisher Scientific). Extracted samples were assessed for RNA quality using the TapeStation 2200 and High Sensitivity RNA reagents (Aligent Genomics, Integrated Sciences). The 10 samples with the highest concentration were pooled using equal concentrations and subsequently concentrated using the RNeasy MinElute Cleanup Kit (Qiagen) (Table S1).

Libraries were constructed using the TruSeq total RNA library preparation protocol (Illumina) and rRNA was removed using the Ribo-Zero-Gold kit (Illumina). Paired-end sequencing (100bp) of the RNA library was performed on the HiSeq2500 platform. All library preparation and sequencing was carried out at the Australian Genome Research Facility (AGRF, Melbourne). All reads have been deposited in the Sequence Read Archive (SRA; BioProject XXX).

### RNA virus discovery

Sequence reads were demultiplexed and trimmed with Trimmomatic followed by *de novo* assembly using Trinity [28]. No filtering of host/bacterial reads was performed before assembly. All assembled contigs were compared to the entire non-redundant nucleotide (nt) and protein (nr) database using blastn and diamond blast [29], respectively, setting an e-value threshold of 1×10^−10^ to remove potential false-positives.

Abundance estimates for all contigs were determined using the RSEM algorithm implemented in Trinity. All viral contigs that returned blast hits with paired abundance estimates were filtered to differentiate those with invertebrate [30], lower vertebrate [31], plant or bacterial host associations using the Virus-Host database (http://www.genome.jp/virushostdb/). The list was further cross-referenced against a known list of viral contaminants [32].

### Virus genome annotation and phylogenetic analysis

Contigs greater than 1000bp in length were inspected using Geneious R10 (Biomatters, New Zealand), and open reading frames (ORF) corresponding to predicted genome architectures based on the closest reference genomes were interrogated using the conserved domain database (CDD, https://www.ncbi.nlm.nih.gov/Structure/cdd/cdd.shtml) with an expected e-value threshold of 1×10^−5^. Reads were subsequently mapped back to viral contigs using bowtie2 [33]. Viruses with full-length genomes, or incomplete genomes possessing the full-length RNA-dependant RNA polymerase (RdRp) gene, were used for phylogenetic analysis. Briefly, amino acid sequences of the polyprotein or gene encoding for the RdRp were aligned using MAFFT [34], and gaps and ambiguously aligned regions were stripped using trimAL [35]. The best-fit model of amino acid substitution was then determined for each data set, and maximum likelihood trees were estimated using IQ-TREE [36]. In the case of coronaviruses, avastroviruses, and avian avulavirus, phylogenies were also estimated using the nucleotide sequences of complete or partial reference genome sequences to better place viruses in context of currently described avian viral diversity. Novel viral species were identified as those that had <90% RdRp protein identity, or <80% genome identity to previously described viruses. Newly identified viruses were named after shipwrecks surrounding King Island, Tasmania. Genome sequences have been deposited in NCBI GenBank (accession numbers XX-XX).

### Viral diversity and abundance across libraries

Relative virus abundance was estimated as the proportion of the total viral reads in each library (excluding rRNA). All ecological measures were estimated using the data set comprising viruses associated with birds and mammals, albeit with all retroviruses and retrovirus-like elements removed (hereafter, “avian virus data set). Analyses were performed using R v 3.4.3 integrated into RStudio v 1.0.143, and plotted using *ggplot2*.

Both the observed virome richness and Shannon effective (alpha diversity) were calculated for each library at the virus family and genus levels using modified Rhea script sets [37], and compared between avian orders using t-tests. Beta diversity was calculated using the Bray Curtis dissimilarity matrix and virome structure was plotted as a function of nonmetric multidimensional scaling (NMDS) ordination and tested using Adonis tests using the *vegan* [38] and *phyloseq* packages [39].

To mitigate the reporting of false-positives due to metagenomic index-hopping, contigs were assumed to be the result of contamination from another library if the read count representing the abundance was less than 0.1% of that representing the highest count for that virus among the other libraries. Short, low abundance contigs from viral genera not detected in any other library sequenced on the same lane were retained.

## Results

### Overall virus abundance and diversity

We characterized the total transcriptomes of Ruddy Turnstones from different age categories upon arrival and prior to departure of migration (Fig 1, Table S1). All individuals were captured in March/November, regardless of age category, and on the same beaches on King Island, Tasmania.

There was a large range in total viral abundance (0.16– 9.91% viral reads) and putative avian viral abundance (0.0071– 1.13% viral reads) in each library (Table S1, Fig. S1). In addition to viral reads that were likely associated with *bona fide* avian viruses, libraries had numerous reads matching invertebrate, plant and bacterial viruses as well as retroviruses (Fig. S1). Although these retroviruses are likely associated with birds, the challenge of differentiating between endogenous and exogenous retroviruses meant that they were excluded from the analysis, as were those viruses most likely associated with arthropods, plants, and bacteria. In total, 11 of the 14 viruses identified in this study likely represent novel avian viral species (Table S2, Fig. 2). Novel species were identified in the double-stranded RNA viruses (*Reoviridae*, genus *Rotavirus* and genus *Orthoreovirus*), positive-sense single-stranded RNA viruses (*Astroviridae*, *Picornaviridae* genera *Hepatovirus*, *Gallivirus*, and unassigned genera), and negative-sense single-stranded RNA virus which is potentially an arbovirus (*Bunyavirales*, *Phenuiviridae*).

**Figure 2.**
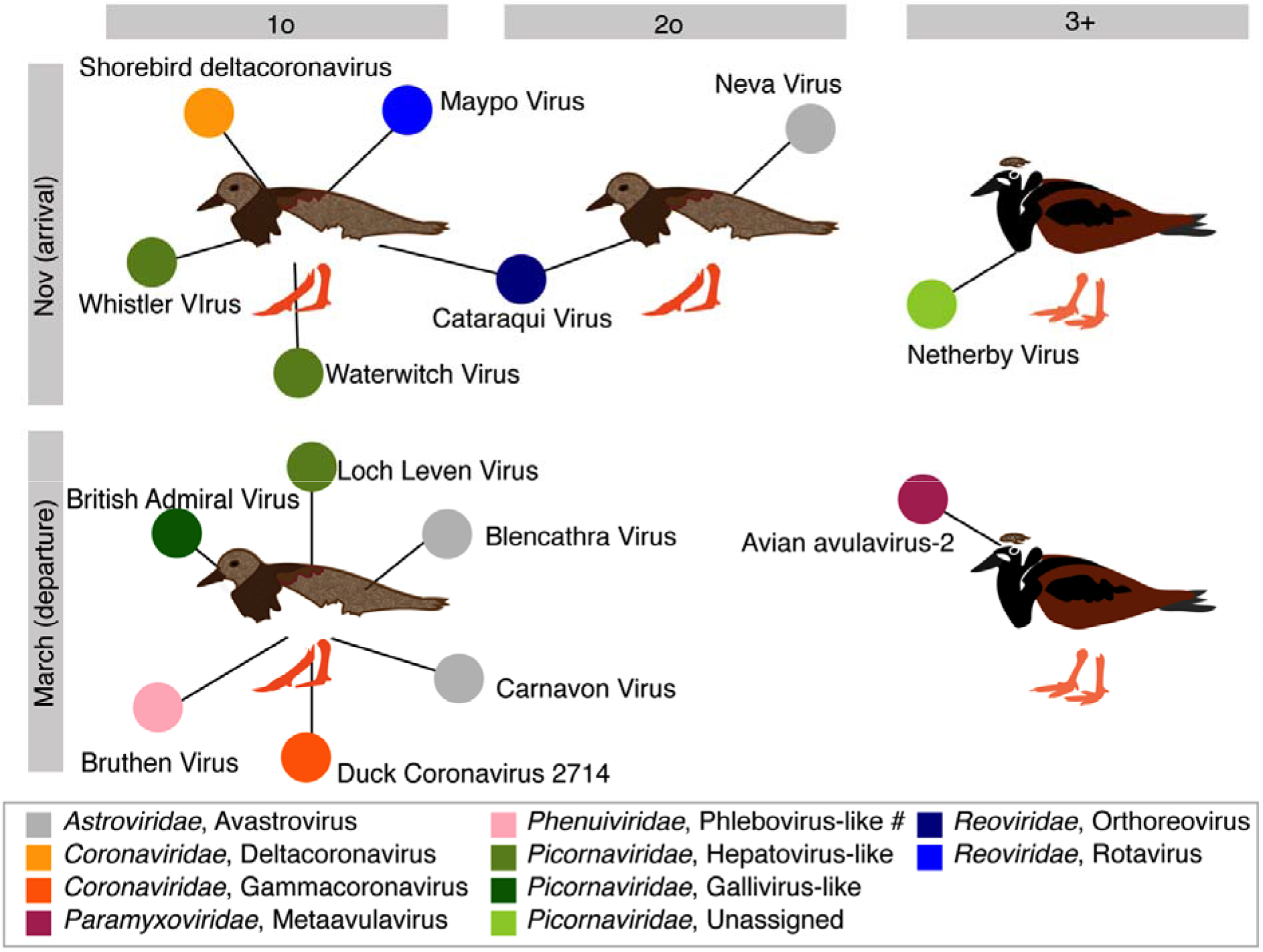
Bipartite network illustrating species for which complete viral genomes were found in each library, comprising 10 individuals. Each library is represented as a central node, with a pictogram of the respective age cohort of Ruddy Turnstone, surrounded by each viral species. Where two libraries share a virus species, the networks between the two libraries are linked. Placement of libraries is arranged by sampling date on the y-axis and age on the x-axis. Virus colour corresponds to virus taxonomy. A list of viruses from each library is presented in Table S2, and phylogenetic trees for each virus family and species can be found in Fig 3–5, Fig S2–S7.

### Novel double-stranded RNA viruses

We identified two novel viruses in two genera of the family *Reoviridae* - *Rotavirus* and *Orthoreovirus*. Maypo virus, a *Rotavirus*, was at very low abundance (0.01% reads) and no full segments were identified with the exception of VP7 that was used for classification within rotavirus species. Maypo virus was separated by a long branch from a clade containing rotavirus G, B, I and viruses identified previously in metagenomic studies (Fig S2). Cataraqui virus, an *Orthoreovirus*, fell as a sister lineage to a recently described avian orthoreovirus - Tvarminne virus (Fig 3A). Cataraqui virus was detected in two libraries - juvenile (1o) and adolescent (2o) birds sampled concurrently - with similar abundance levels (0.5% and 0.25% of reads in each library, respectively) showing viral sharing across these age groups (Fig 3A, Fig 6). The two genomes recovered were highly similar, with only ~1 nucleotide difference in each of the lambda segments. While we could not assemble this genome from adult 3+ birds, contigs of Cataraqui virus were also found, comprising 0.0023% of the library. As this represents >0.1% of the read abundance of the library in which this virus is at its highest abundance we tentatively exclude index-hopping.

**Figure 3.**
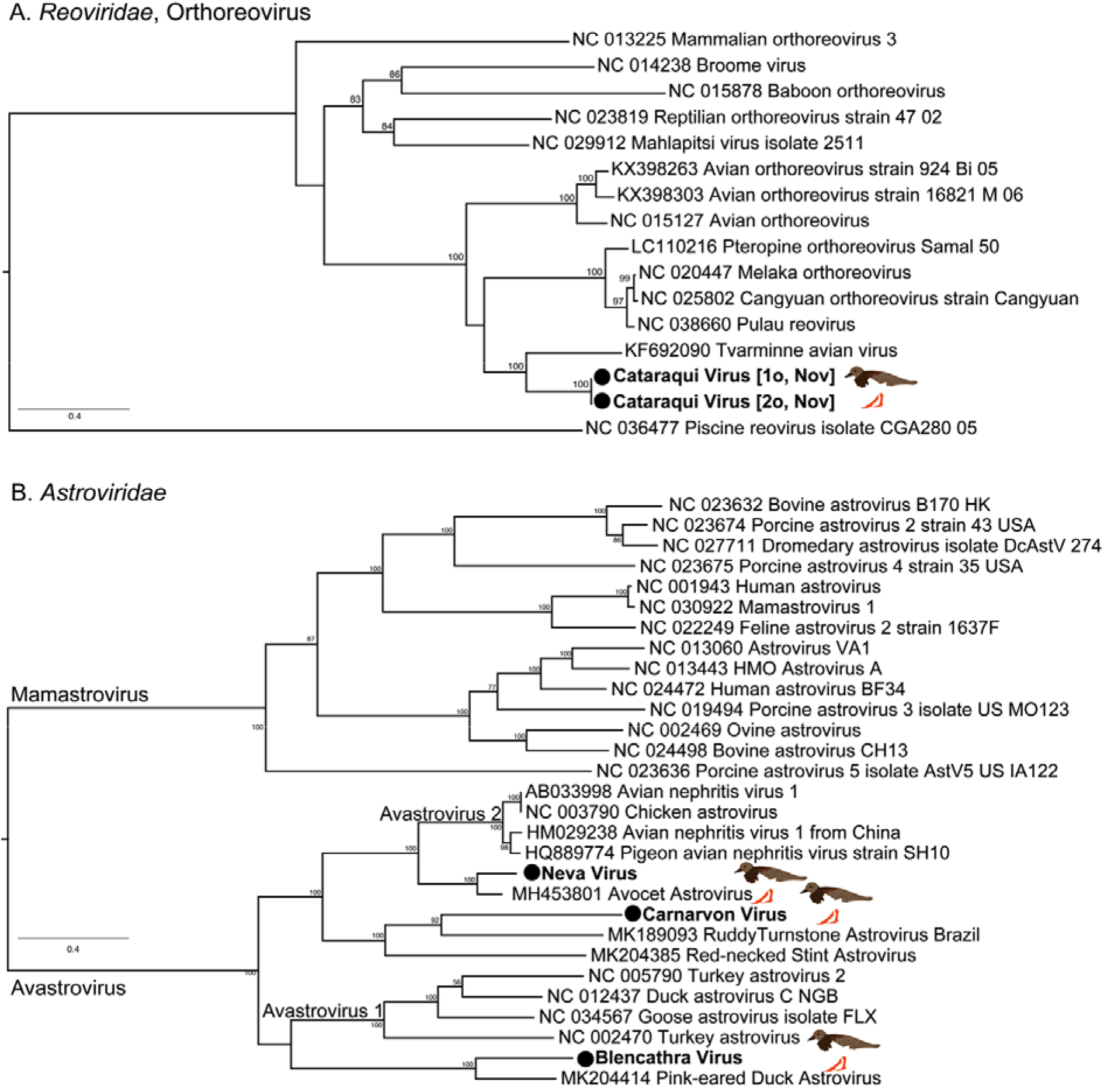
(A) Phylogenetic tree of the LambdaB segment, containing the RdRp, of orthoreoviruses. Piscine reovirus is set as the outgroup. (B) Phylogenetic tree of the ORF1ab, including the RdRp, of avastroviruses. Avastrovirus 3 is not shown as there are no full genomes available, but is presented in Fig S2 which shows a phylogeny inferred from a short region of the RdRp. The tree was midpoint rooted, corresponding to division between mammalian and avian viruses. The sequences generated in this study are indicated by a filled circle and are in bold. Bootstrap values >70% are shown for key nodes. The scale bar indicates the number of amino acid substitutions per site.

### Novel single-stranded RNA viruses

Three novel avastroviruses were identified in this analysis, two in the library of 1o birds sampled in March and one in the 2o birds sampled in November. Specifically, although Neva virus and Carnarvon virus occupy different phylogenetic positions, both are related to viruses previously classified as avastrovirus 2 (which includes avian nephritis virus), while Blencathra virus is a sister lineage to an astrovirus recently described in the pink-eared duck [40] (Fig 3B, Fig S3).

We also identified five picornaviruses (family *Picornavirdae*). Three of these - Waterwitch virus, Whister virus, and Loch Leven virus – fall as sister lineages to members of the genus *Hepatovirus* and *Tremovirus* and were found exclusively in 1o birds, with Waterwitch virus and Whister virus forming a distinct clade (Fig S4). In contrast, British Admiral virus fell as a divergent sister-group to the *Gallivirus* genera, while Netherby virus, the only picornavirus found in a 3+ bird, fell into a currently unassigned clade comprising a chicken and a quail phacovirus (Fig S4).

We also identified a novel virus in the family *Phenuiviridae*, Bruthen virus. This virus fell in a group of unclassified viruses that are the sister-group to members of the genus *Phlebovirus* that contains numerous tick-borne viruses (Fig 4). In the phylogeny Bruthen virus is most closely related to Laurel Lake virus sampled from *Ixodes scapularis* ticks in the USA [41]. However, because of the long branch lengths across the tree as a whole, including that leading to Bruthen virus, it is currently unclear whether this virus is a strictly arthropod virus or an arthropod-borne virus that may infect birds (Fig 4).

**Figure 4.**
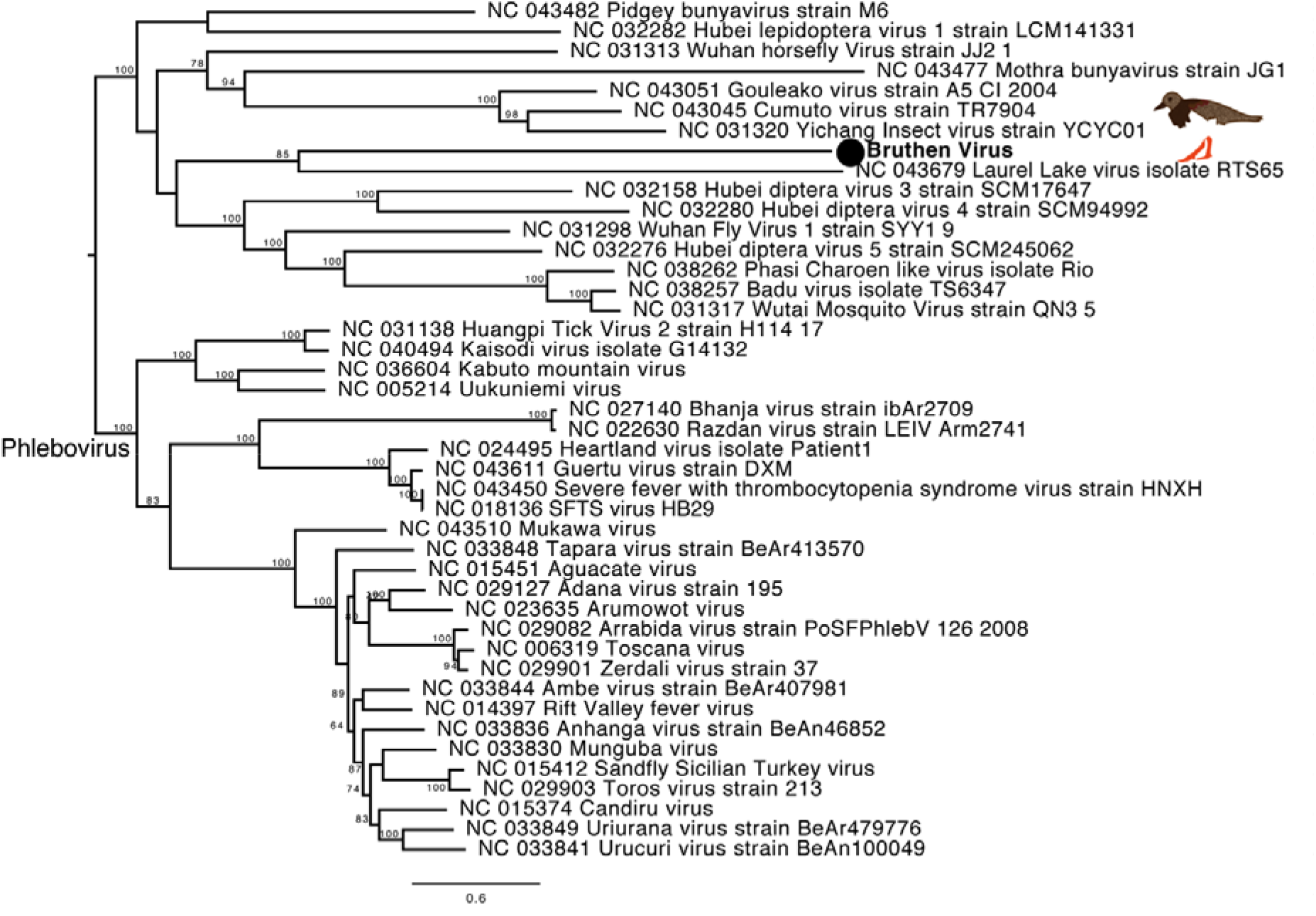
Phylogeny of the polyprotein of the *Phenuiviridae*. The tree was midpoint rooted for clarity only. Viruses described in this study are marked in bold, adjacent to a filled circle. Bootstrap values >70% are shown for key nodes. The scale bar indicates the number of amino acid substitutions per site.

### Identification of previously described avian viruses

We identified two members of the *Coronaviridae* – Duck coronavirus 2714 (genus *Gammacoronavirus*) and a virus here referred to as Shorebird deltacoronavirus (genus *Deltacoronavirus*) (Fig 5, Fig S5-S6). These viruses are common in wild waterbirds, although most sequence data is limited to short fragment of the RdRp (Fig S8, S9) [42]. In the ORF1ab phylogeny, the Duck coronavirus 2714 sequence (subgenus *Igacovirus*) was most closely related to a sequence identified in gulls in Canada [43], however based on a phylogeny comprising 400bp of the RdRp, the sequence revealed here fell in a clade dominated by viruses previously identified in Australia, and most closely related to viruses identified in Ruddy Turnstones from King Island, sampled in 2016 (Fig 5, Fig S5) [44]. In the ORF1ab phylogeny the Shorebird deltacoronavirus fell as a sister lineage to the subgenus *Buldacovirus*. Based on a phylogeny comprising 400bp of the RdRp, for which more sequences are available, this virus was most closely related to viruses previously identified in Australia, including shorebirds from Victoria and Ruddy Turnstones from King Island in 2016 (Fig S6) [44]. The genome of Shorebird deltacoronavirus revealed here comprises the first full genome of this putative viral species, and the detection of two different coronaviruses from these birds suggests Ruddy Turnstones may be important reservoirs for both gamma and deltacoronaviruses.

**Figure 5.**
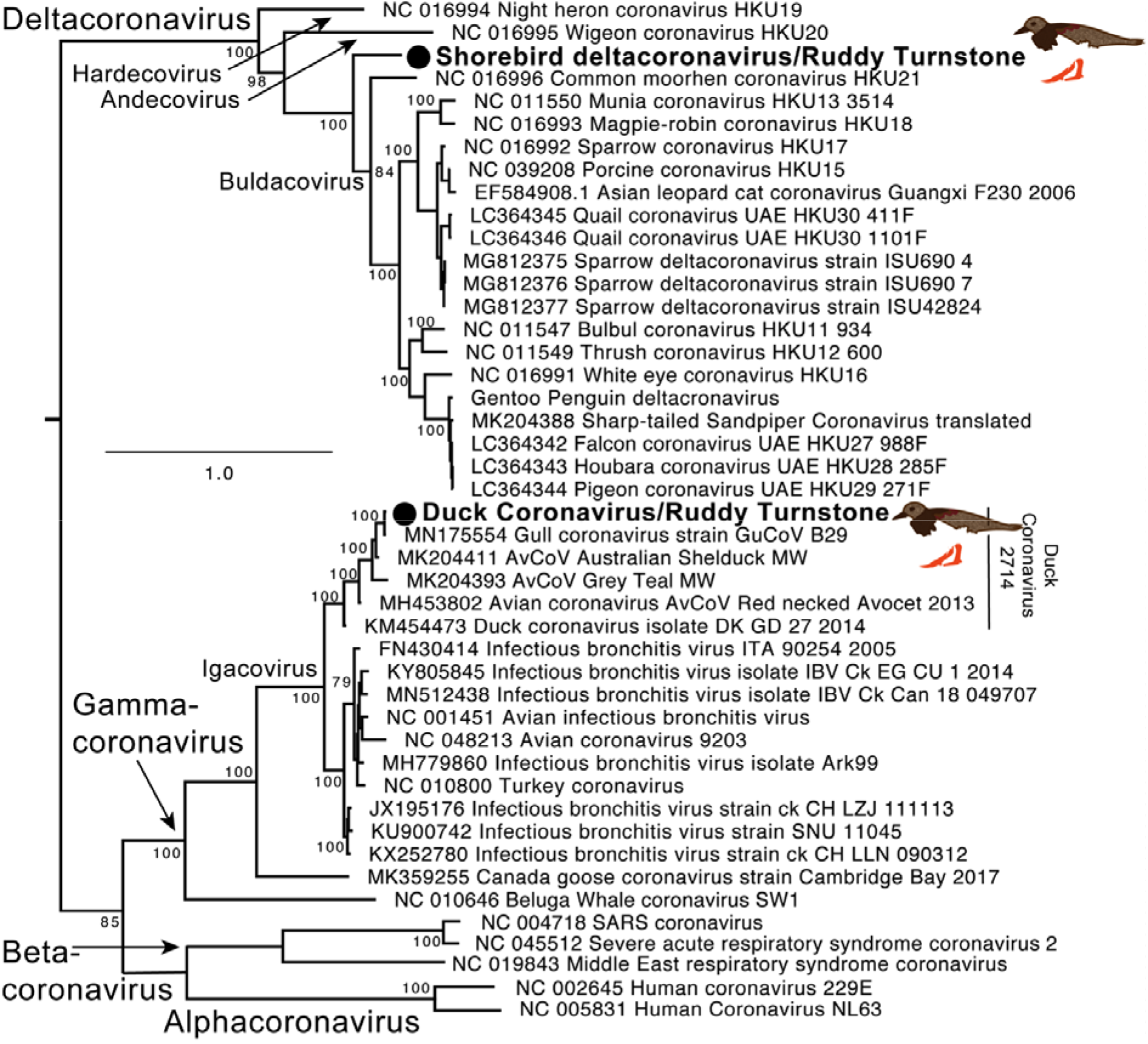
Phylogenetic tree of the ORF1ab, containing the RdRp, of the *Coronaviridae*. The sequences generated in this study are indicated by a filled circle and are in bold. Deltacoronaviruses are set as the outgroup, as per [42]. Only select members of the alpha- and betacoronaviruses are included. Bootstrap values >70% are shown for key nodes. The scale bar indicates the number of amino acid substitutions per site. Partial gammacoronavirus and deltacoronavirus phylogenies, including numerous wild bird sequences, are presented in Fig S5 and S6, respectively.

The variant of avian avulavirus 2 identified here (AAvV-2/Ruddy turnstone/Tasmania/2015) was a (weakly supported) sister-group to a clade of viruses described in wild birds in China (Fig S7), and we speculate that this likely represents a lineage that is more common in wild birds than poultry. Unlike the case of avian avulavirus 1 (Newcastle Disease), the polybasic cleavage site within the F gene is not a determinant of the pathogenicity of avian avulavirus-2in chickens [45, 46]: as such, we have not performed cleavage site annotation. This virus had a very high abundance in the library comprising adult (3+) birds sampled in March (comprising 99% of all avian viral reads and 0.2% of all reads). Reads for this virus were also identified in the juvenile (1o) birds simultaneously sampled, and adolescent (2o) birds sampled in November, although with very low abundance (0.00079% reads and 0.00014% reads, respectively, but above the assumed index-hopping threshold of 0.1% of the most abundant virus).

### Age affects both virome diversity and abundance

Regardless of sampling period, juvenile (1o) birds had higher viral abundance and diversity than adult (3+) birds at the viral family, genus and species levels. First, viral abundance was higher in juvenile than adult birds, with a 10-fold difference in November (1.12% compared to 0.007% avian viral reads) and March (0.2% compared to 0.02% avian viral reads). Adolescent (2o) birds, which overwintered on the island rather than making a return trip to the Arctic breeding grounds like the 3+ birds did, also had very high viral abundance in November (0.7% avian viral reads) (Fig 6A). Second, this trend remained when assessing diversity within each sample (i.e. alpha diversity). Observed richness and Shannon diversity was higher in juvenile than adult birds, in which observed richness was significantly higher in 1o birds at the viral genus level (Fig S8) (t = 5.8138, df = 1.4706, p= 0.049). At the viral species level, there was a significant difference in the number of viral species in juvenile compared to adult birds (t = 9, df = 1, p= 0.05): in November and March we identified five and six viruses in juvenile birds, respectively, compared to one virus from each of the libraries comprising adult birds, with adolescent birds exhibiting intermediate values (two virus species described) (Fig 2, Fig 6). Although this study is clearly limited in sample size, the overall trend remains across our biological replications (here different time periods). Sampling period was not a significant predictor of either observed richness or Shannon diversity at the virus family, genus and species levels. For example, when assessing the number of viral species, we observed the same number of viruses revealed (excluding the adolescent 2o birds as they were only sampled in November) in both November (n=6) and March (n=7) (Fig 2).

**Figure 6.**
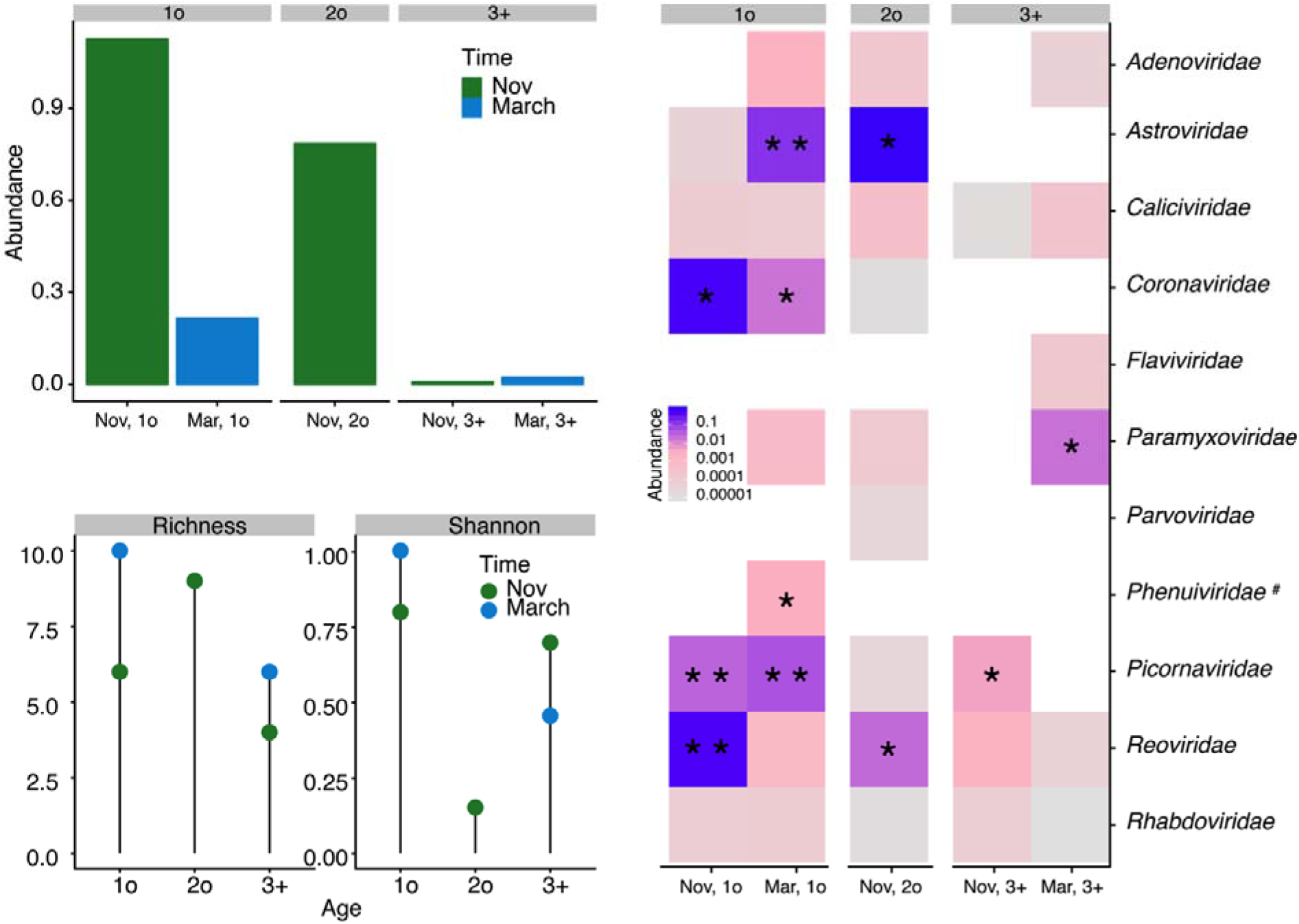
Effect of age on abundance and alpha diversity. (A) Viral abundance is higher in 1o birds compared to 3+ birds. (B) Alpha diversity at the viral family level, here measured as observed richness and Shannon diversity, is higher in 1o birds compared to 3+ birds. Alpha diversity at the virus genus level is presented in Fig S1. (C) Heatmap illustrating viral diversity, at the genus level in each library, with colour corresponding to viral abundance. Blue and purple correspond to viruses with high abundance, and pink corresponds to viruses with low abundance. Asterisks indicate cases in which at least one complete viral genome was obtained. Viral families marked with a hash symbol indicate families where host association is unclear. Plots for the virus genus level are presented in Fig S8.

To ensure this trend was not due to a sequencing artefact we tested for differences in viral abundance and alpha diversity in those viruses sampled from birds but that most likely infected invertebrate hosts. These invertebrate viruses effectively comprise a null model: because they do not actively replicate in birds we would expect to see similar abundance levels in both the juvenile and adult birds. As expected, there was no statistical difference between juvenile and adult birds (t = 1.0022, p = 0.5). Furthermore, juvenile and adult birds exhibited similar levels of both alpha diversity measures for invertebrate viruses (observed richness or Shannon diversity), regardless of whether the measure was performed at the virus “cluster” level [30] (Observed Richness t = −1.3416, p = 0.3499; Shannon Diversity t = −0.6925, p = 0.5606) or at the virus family level (and family-like when no family classification was available) (Observed Richness t = 0.56569, p = 0.6693; Shannon Diversity t = −0.54246, p = 0.6425) (Fig S9).

No obvious trends were observed with respect to beta diversity (Fig S10). Specifically, neither age (R^2^=0.49, p=0.33) nor sampling time point (R^2^=0.2, p=1) was associated with avian virome structure at the viral genus or family levels. There was very limited viral sharing (i.e. connectivity) among birds of different age categories even though they were sampled on the same beaches at the same time. The only viral species that we were able to annotate from more than one library was Cataraqui virus, identified in juvenile and adolescent birds at the November time point: however, reads for this virus were also present in the adult birds in November (Fig 2, Fig 3A). While the full genome of avian avulavirus-2 was only identified in the adult birds in March, reads were present in juvenile birds from March. Despite this, due to cohabitation, virus connectivity was far lower than expected. We suggest that adult birds may already have antibodies against many of these viruses, resulting in no detectable infections in this age category.

## Discussion

We used a meta-transcriptomic approach to help reveal the role of host age in shaping the avian virome. Through this, we identified 14 viruses, of which 11 are novel, in apparently healthy wild migratory birds from a single sampling location in Australia. The largest diversity of viruses were from the families *Astroviridae*, *Coronavirdae*, *Picornaviridae* and *Reoviridae*, from which more than one putative virus species was identified in each case.

Based on studies of the ecology of both avian influenza A virus [1, 7, 8], and more recently, of virus communities in bats [47], we predicted that juvenile birds would have high viral abundance and diversity compared to adult birds. During the non-breeding period, Ruddy Turnstones on King Island, Tasmania spend time in small (~50 birds) groups comprising both adult and juvenile birds, and are often restricted to small stretches of coastline (~1 km). Importantly, our sampling collection and selection strategy means that within each sampling period (November, and March) we compare birds of different age groups that are cohabiting, enabling us to identify any viruses that are shared between age groups within a given sampling period. Indeed, we observed high viral abundance and diversity in juvenile birds compared to very low abundance and diversity in adult birds. Adolescent birds shared a viral species, Cataraqui Virus, with co-sampled juvenile birds, demonstrating a degree of viral sharing among these two age categories. The high viral diversity and abundance in juvenile birds, but low levels in conspecific adult birds, strongly suggests that adult birds have immunity against these viruses. As a consequence, our results raise important questions about the robustness and longevity of antibodies against viruses in wild birds, such as the Ruddy Turnstone. Work on avian influenza A viruses suggests that short-lived species, such as dabbling ducks, may have short lived antibodies [48] and may therefore be reinfected many times [49], in comparison to long lived species such as swans [15] and shearwaters [50] that had detectable, neutralizing antibodies for years. Our data suggest that Ruddy Turnstones may fit into the latter category, although this needs to be clarified by antibodies studies.

Due to limited sample size, we utilized a biological replicate to test the impact of age. Specifically, we sampled birds following their arrival on King Island and prior to their departure following their non-breeding staging period. Despite only being four months apart, we found no evidence of viral connectivity between these two time points. This clearly demonstrates the transient nature of the avian virome, in which there was no consistent detection of viral species across sampling points. Further, we found no difference in viral abundance or diversity of adult or juvenile birds when comparing those captured in November (recently arrived) or the subsequent March (imminently prior to departure of the 3+ adult birds). This not only shows the lack of a migratory cycle effect, but also that the impact of age is strong, despite potentially large differences in physiology of Ruddy Turnstones at these sampling events. Prior to migration, birds go through a period of impressive mass increases; some species may more than double their body mass, with a 1-3% increase per day [51, 52]. Indeed, birds captured in March had an additional 50% in mass compared to November. Studies in the microbiome of migratory birds found that the majority of bacterial taxa are not affected by migration when comparing resident birds and those under active migration immediately upon arrival [53]. Other changes during active migration may be related to immune responses. It has been suggested that migrating animals should boost their immune function during migration [54, 55]. A wind tunnel experiment found no difference in immune function between “migrating” Red Knots (*Calidris canutus*) and control birds [56]. In contrast, migratory Common Blackbird *Turdus merula* individuals had a lower innate immune function during migration compared with resident individuals [57]. Despite the different migration dispositions between the November and February time points, we found a consistent age effect and no difference in abundance, alpha diversity or beta diversity across time points. We also confirmed that this age effect was not an artefact by using invertebrate-associated viruses, that do not replicate in birds, as an intrinsic biological control. Importantly, we found that invertebrate associated viruses demonstrated no differences in viral abundance or alpha diversity in juvenile compared to adult birds, in marked contrast to the pattern in ‘true’ avian viruses. This observation is consistent with previous studies of virus ecology, in which influenza A viruses exhibit higher prevalence in juveniles compared to adults [7, 8], and juveniles have a greater number of infections and diversity of subtypes over an autumn season compared to adult birds [22]. Similarly, caves hosting a larger proportion of juvenile bats had a larger viral diversity as compared to those hosting a low proportion of juvenile bats [47].

Our results allow us to reflect on the potential barriers to migration, specifically why some individuals remain on non-breeding areas and do not migrate. Annual migrations are undertaken by adult shorebirds, that also have with lower viral abundance and diversity. We can therefore hypothesize that one of the reasons adolescent birds do not undertake a northward migration in their second year may be increased virus susceptibility, itself due to lower age-dependant immunity. Here, we show high viral abundance in both juvenile and adolescent birds. Even without obvious disease, the infection status and intensity of some viral infections may have negative effects on body stores, refuelling capacity, movement, phenology and survival [58]. These effects may result in “migratory culling” - the combined physiological effects of migration and infection mitigation that may remove individuals from the population [59]. This hypothesis may be central to understanding an important barrier to migration of juvenile and adolescent birds, given demonstratively high viral abundance and diversity compared to adult conspecifics, and clearly merits further study.

Overall, we demonstrate a large viral diversity in Ruddy Turnstones and markedly different viromes that were detected in birds across age groups and between sampling periods. This, in turn, highlights the transient nature of the avian RNA virome and the snap-shot nature of virome studies to date. Both these characteristics have recently been observed in studies of bat viromes [47], strongly suggesting this is an important consideration in all ecological studies that utilise virome data. Beyond ecological studies, this is a critical consideration for future surveillance efforts for novel and emerging viruses. Finally, we further demonstrate that apparently healthy wild birds are able to sustain high viral loads and diversity without obvious signs of disease. This, combined with the continued detection of viral families previously associated with diseases such as gastroenteritis or hepatitis [60, 61], suggests that many of the viruses may not be pathogens.

## Acknowledgements

The sampling of birds in Australia importantly relied on the assistance of volunteers within the Victorian Wader Study Group, particularly Clive Minton, Robyn Atkinson and Rob Patrick; and members of the Centre for Integrative Ecology at Deakin University, notably Simeon Lisovski and Bethany Hoye. The sampling was supported by NIH/NIAID (HHSN2662007 00010C), ARC discovery grants (DP 130101935 and DP160102146) and an ARC Australian Laureate Fellowship to ECH (FL170100022). MW is funded by an ARC Discovery Early Career Researcher Award (DE200100977).

